# Near-infrared spectroscopy discriminates mass-reared sterile and wild tsetse flies

**DOI:** 10.1101/2024.08.26.609677

**Authors:** Soumaïla Pagabeleguem, Dari F. Da, Bernard M. Somé, Marx S.P. Avelessi, Nicaise D. C. Djègbè, Rebecca L. Yoda, Abdramane Bagayogo, Hamidou Maïga, Thomas S. Churcher, Roch K. Dabiré

**Author notes:** Co-lead authors (Soumaïla Pagabeleguem, Dari F. Da). **E-mail addresses:** BMS MSPA NDCD RLY AB HM TSC RKD.

## Abstract

In the frame of the Pan African Tsetse and Trypanosomosis Eradication Campaign, an integrated pest management approach has been adopted to manage tsetse fly populations using a sterile insect technique (SIT).. Monitoring the efficacy of the SIT programs requires the discrimination between wild and sterile males collected in monitoring traps. The discrimination between sterile and wild males relies mainly on marking of sterile males with a fluorescent dye powder (before the releases) and their identification using a fluorescence camera and /or a fluorescence microscope. However, the accuracy of this method remains limited with defective marking and wild flies contaminated with a few dye particles in the monitoring traps. The molecular techniques developed to discriminate doubtful flies are costly for endemic countries. Here, we investigate the ability of a new generation monitoring tool, the near-infrared spectroscopy (NIRS) to discriminate between lab-reared *Glossina palpalis gambiensis* and their counterparts in the field. NIRS technique discriminates wild flies males with 86% accuracy to *Glossina palpalis gambiensis* males reared in an insectarium. Interestingly, the prediction accuracy between wild and laboratory-reared flies improved to 88% when the *Glossina* lab colony flies were irradiated. In light of these results, NIRS can discriminate tsetse flies even when identification with the UV camera is doubtful. However, more investigations integrating some natural variables are required before the validation of the technical as a complementary method for potential future tsetse eradication programs.

## Introduction

Tsetse flies (*Glossina spp*.) are the only cyclic vectors of trypanosomes, responsible for sleeping sickness or Human African Trypanosomosis (HAT) in humans and nagana or African Animal Trypanosomosis (AAT) in livestock in sub-Saharan Africa (Vreysen et al. 2013). Tsetse infests 38 countries in Sub-Saharan Africa, hindering the development of sustainable and productive agricultural systems over more than 10 million km^2^ (Hursey et Slingenbergh 1995). Over 60 million people are continuously exposed to the risk of trypanosomiasis infection, and farmers in tsetse-infested areas suffer up to 20-40% losses in livestock productivity leading to an estimated annual loss of about USD 4.75 billion (Swallow 1999).

In order, to eliminate or eradicate these vector-borne diseases, four methods that are environmentally and economically acceptable, can be used in the context of area-wide integrated pest management (AW-IPM) approaches. These strategies include the sequential aerosol technique, the deployment of insecticide-impregnated traps/targets, the pour-on, and the sterile insect technique (SIT) (Vreysen et al. 2013; Bouyer et al. 2013). AW-IPM program with an SIT component has been used throughout the world to suppress, eradicate, contain or prevent the introduction of several insect pests such as fruit flies (Hendrichs et al. 2021), moths (Bloem et al. 2007), screwworm flies (Vargas-Terán et al. 1994), mosquitoes (Klassen et Curtis 2005) and tsetse (Vreysen et al. 2000). Furthermore, the effectiveness of the SIT has been demonstrated for the eradication of tsetse populations in numerous African countries (Zanzibar, Nigeria, Burkina Faso) where different vectors have been highlighted (Vreysen et al., 2000, Takken et al. 1986; (Cuisance et al. 1984; Politzar et Cuisance 1984). In Senegal, a program is underway to eradicate a *G. p. gambiensis* population from the Niayes area (Pagabeleguem et al. 2015; Seck et al. 2024; Vreysen et al. 2021).

In AW-IPM programs with SIT component, the status of the field-trapped flies (mass-reared sterile males or wild fertile males) remains one of the most crucial prerequisites for success. Indeed, the ratio of sterile over wild flies estimated is an important parameter for monitoring the efficiency of SIT campaign (Vreysen et al. 2000; Vreysen 2005; Vreysen et al. 2014). The discrimination between sterile and wild males relies mainly on the fluorescence marking of insectary reared flies. The fluorescence marking is incorporated during the adult’s emergence using a fluorescent dye powder (DayGlo®) mixed with sand or after emergence by impregnation with the powder. This treatment makes a discrimination of sterile from wild flies using a fluorescence camera and /or a fluorescence microscope (Pagabeleguem et al. 2015).

However, marking sterile males with fluorescent dye-powder has some limits. Indeed, the fluorescent dye powder is not infallible, and some sterile flies could be poorly marked with very few powder particles. Conversely, wild flies could be contaminated in the monitoring traps with some powder particles for being in contact with their sterile counterparts. Flies could also be predated by ants in the traps and then lose their head, which makes difficult the accurate determination of their origin. To overcome this challenge, a molecular tool has been developed allowing discrimination with high accuracy between the mass-reared sterile males and their wild counterparts (Pagabeleguem et al. 2016). However, this new method is costly and requires a significant amount of time to anayse a large number of samples results (Pagabeleguem et al. 2016). Consequently, it is urgent to develop alternative tools that could be not only accurate but also rapid, and cost-effective for the discrimination of sterile and wild males.

The Near-infrared spectroscopy (NIRS) technique, a potential new-generation monitoring tool, could be a reliable alternative. The NIRS technique principle measures the amount of near-infrared energy absorbed by biological samples at specific wavelengths. This method is rapid, non-destructive, cost-effective, reagent-free and requires little technical training of personnel following algorithm development (Sikulu-Lord et al. 2024; Goh et al. 2021; Mayagaya et al. 2009). In previous investigations, NIRS has been explored in malaria area for predicting mosquito age, *Anopheles* species and theirs status for *Plasmodium* infection (Sikulu et al. 2010; Mayagaya et al. 2009; Da et al. 2021; Maia et al. 2019), triatomine species (Depickère, Ravelo-García, et Lardeux 2020) and *Biomphalaria* species (Valladares et al. 2021) and to separate the sex tsetse pupae (Argilés-Herrero et al. 2023; Dowell et al. 2005). It has also been explored to detect the *Trypanosoma cruzi* in midgut, rectum and excreta samples of *Triatoma infestans* (Tátila-Ferreira et al. 2021). This study aimed to determine whether the NIRS technique could accurately discriminate between wild caught and lab-reared, sterile and markedmale *Glossina palpalis gambiensis*. .

## Methods

### Study design

The empirical method used to discriminate lab sterile *Glossina* from wild flies is based on detecting fluorescent powder marking in the laboratory reared-flies using a fluorescence camera and /or a fluorescence microscope. This study is designed to understand whether NIRS is also able to distinguish unmarked-sterile tsetse males and theirs homologous caught in the field. To achieve this, different analyses were performed to check whether the technique is able to make distinction between wild flies and the laboratory-reared tsetse flies without fluorescent marking, between and also if the irradiation and marking actions in the laboratory can impact the NIRS prediction.

### Insectarium flies

*Glossina palpalis gambiensis* flies used for experiments were from a colony maintained at the Insectarium de Bobo-Dioulasso (IBD) under standard conditions (25 ± 1 °C, 75 ± 5% RH and 12:12 light: dark photoperiod). Historically, the colony was derived from a strain established in 1972 using wild flies collected in Guinguette, a locality close to Bobo-Dioulasso, Burkina Faso (Itard 1976). Then, the resulting colony was mass-reared in the insectarium of the “Centre International de Recherche-Développement sur l’Elevage en zone Subhumide (CIRDES)” (Bauer, et al. 1984). About 54,000 adult flies from CIRDES colony were transferred in 2016 to the IBD insectary as a base of a new colony for mass rearing (Pagabeleguem et al. 2021).

Two experimental groups of insectarium flies were set up: unmarked fertile flies and marked sterile flies. For the unmarked fertile flies, 1000 flies were sampled from new emergence colony flies, divided into Roubaud cages (4.5 × 13 × 8 cm), and placed at standard conditions. For the marked sterile flies, batches of pupae from which almost all females have emerged (about three days after the beginning of emergence) have been chilled at 8°C and irradiated with 120 Gy using a Cobalt 60 source (model 812 S/N 002) (Pagabeleguem et al. 2023). Irradiated pupae were placed in Petri dishes under ∼1 cm of sand mixed with a fluorescent dye (DayGlo®) (0.5 g dye / 200 g of sand) in order to mimic natural emergence conditions and to mark flies during the emergence (Seck et al. 2015). The marking of the sterile male flies is required since during the operational sterile male released program this mark allows discrimination from wild flies in the monitoring traps and to assess SIT program progress (Vreysen 2005). Approximately 500 sterile marked flies (mostly males) were sampled and divided into Roubaud cages, therefore, 20 flies by cage.

Flies of both groups were fed with irradiated bovine blood using *in vitro* silicon membrane system and maintained in the insectary in standard conditions. For NIRS scanning, 60 unmarked and 60 marked flies were sampled every five days from the emergence day until 30 days of age (j1, j5, …. j25 and j30) and shipped immediately to the Institut de Recherche des Sciences de la Santé (IRSS) laboratory for analysis.

### Wild flies

Wild flies were caught in the forest of Bama (11.38454N, 4.409087W), located about 25 kilometers from Bobo-Dioulasso. This forest, a small (500 m long and 130 m large), has permanent water and protected vegetation, as well as some monitor lizards (*Varanus niloticus*), crocodiles (*Crocodylus niloticus*), domestic animals and humans that probably constitute the main feeding source of *G. palpalis gambiensis* population. In September and October 2021, tsetse flies were sampled during two field trips lasting five days each using six biconical traps (Challier et Laveissière 1973) and five monoconical traps (Laveissière et Grébaut 1990). Each of these flies was morphologically identified by sex and species as described by Pollock (1982). Thereafter, they were placed in humidified containers and transferred to the IRSS laboratory where they were immediately scanned.

### Flies scanning

Flies from IBD killed by exposition to chloroform vapor for 3 minutes. Each fly was scanned using a LabSpec4 Standard-Res i (standard resolution, integrated light source) near-infrared spectrometer and a bifurcated reflectance probe mounted 2 mm from a spectral on white reference panel (ASD Inc., Westborough, Massachusetts, USA) (Musiime et al. 2019). All specimens were scanned on the side centering the head and thorax region under the focus of the light probe, and the RS3 spectral acquisition software (ASD Inc., Malvern PANalytical) was used to record the flies’ spectra. This software automatically records the average spectra from 20 scans. Absorbance was recorded from 350 to 2500 nm of the electromagnetic spectrum.

### Data analysis

The statistical analysis was based on the spectra collected from the scanning of laboratory-reared and wild flies. Data analysis was then carried out according to the following procedure: (a) marked sterile flies *versus* wild-caught flies which are the interest group for SIT programs, (b) unmarked sterile flies *versus* marked sterile flies in order to understand the ability of the NIRS to determine the marking status (marked/unmarked) of laboratory-reared flies, (c) unmarked sterile flies *versus* unmarked fertile flies to know whether the NIRS could identify the sterilization status (sterile/fertile) of laboratory-reared tsetse flies, and finally (d) marked sterile flies *versus* wild-caught flies to determine the origin and marking status.

Machine learning methods were used to construct binomial logistic regression models using maximum likelihood. The mean of the two spectra from each tsetse fly was used in the analysis. Spectra were then trimmed to values corresponding to 500 - 2350 nm to remove the excess noise arising from the sensitivity of the spectrometer at the ends of the near-infrared range. These spectra were analyzed using partial least squares regression (PLS), a statistical technique utilizing the covariance between the spectra and marking status in order to extract the most informative elements within a much smaller dimension. All analyses used simple models that did not include spectra smoothing or penalized coefficient function estimation. Data is divided into three subsets (at a 3:1:1 ratio) for model training, validation (where the number of principal components is chosen from 2 to 50), and testing, which estimates the generalized error. All models used random sampling to ensure an equal number of observations per class. This was repeated 100 times, and the models were averaged across each realization. The accuracy of the model was evaluated using the area under the receiver operating characteristic curve value (AUC, value closer to 1 indicating better performance). All analyses were carried out in R using the package mlevcm (Esperança et al. 2020).

## Results

A total of 627 insectary-reared males of *G. palpalis gambiensis* aged from 1 to 30 days and 143 wild-caught tsetse males were used to determine the predictive discrimination ability of the model. Based on the spectra profile, rough differences were perceptible between some experimental groups: The average spectra from field flies *versus* laboratory sterile-unmarked flies differed with globally high absorbance observed in wild *Glossina* (Fig 1).

**Figure 1:**
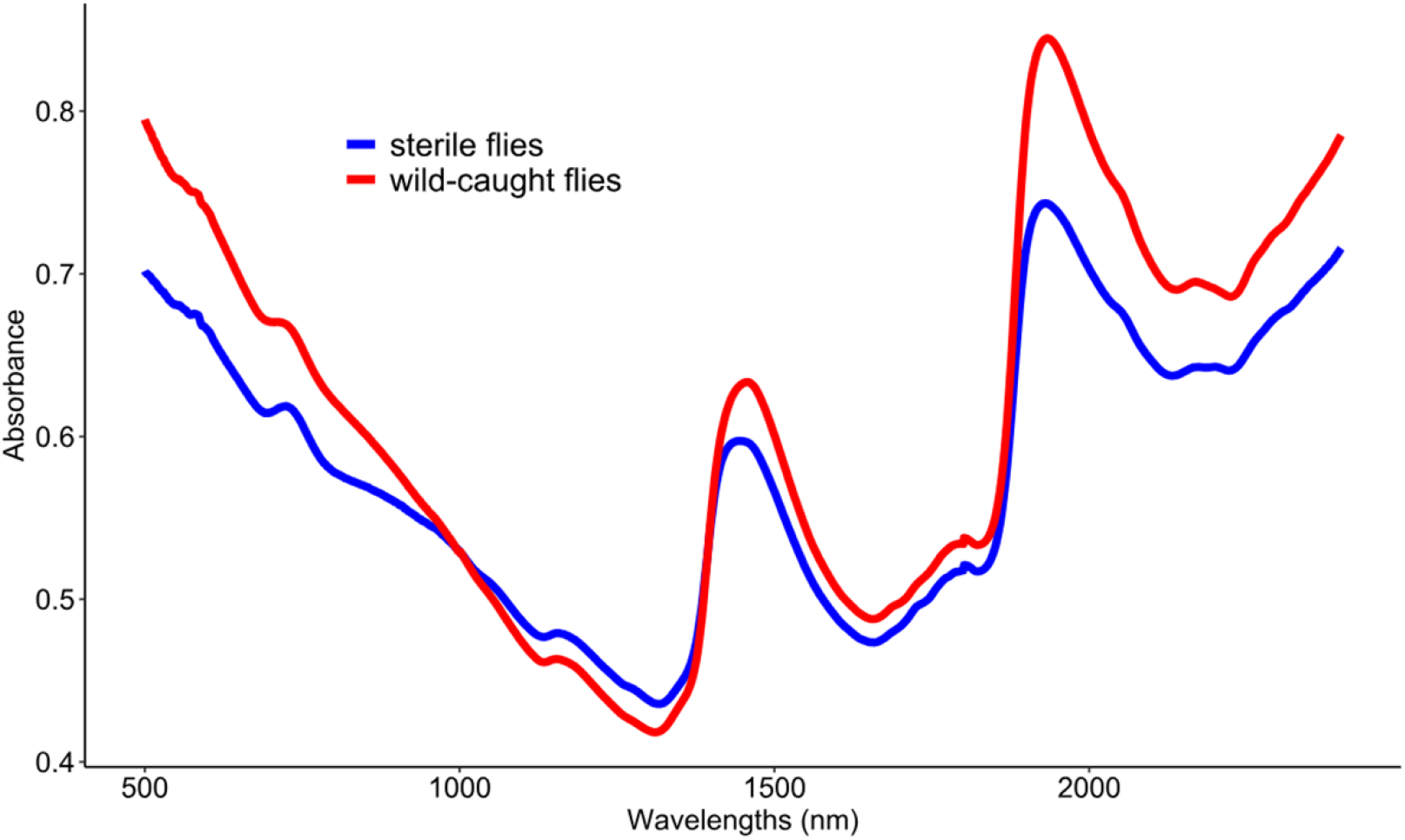
Average spectra of laboratory sterile-unmarked flies and wild-caught flies, each being sampled at discrete wavelengths in the interval [500, 2400 nm].

NIRS demonstrated a remarkable ability to distinguish between marked-sterile males and wild-caught flies with 88% accuracy (marked-sterile: 88%; wild-caught: 87%). More crossed analysis between the groups of *G. palpalis gambiensis* taking into account the interventions operated during flies rearing in the laboratory (marking and irradiating) was done to understand the origin of these observed differences.

For our main question, determining the ability of NIRS to differentiate between sterile and wild tsetse flies (sterile-unmarked vs wild-caught flies), we obtained an overall accuracy of 90% (Fig 2). The same trend was observed when the sterile males were marked with fluorescence. These results suggest that marking with fluorescent powder does not appear to have any impact on the discrimination between sterile and wild flies.

**Figure 2:**
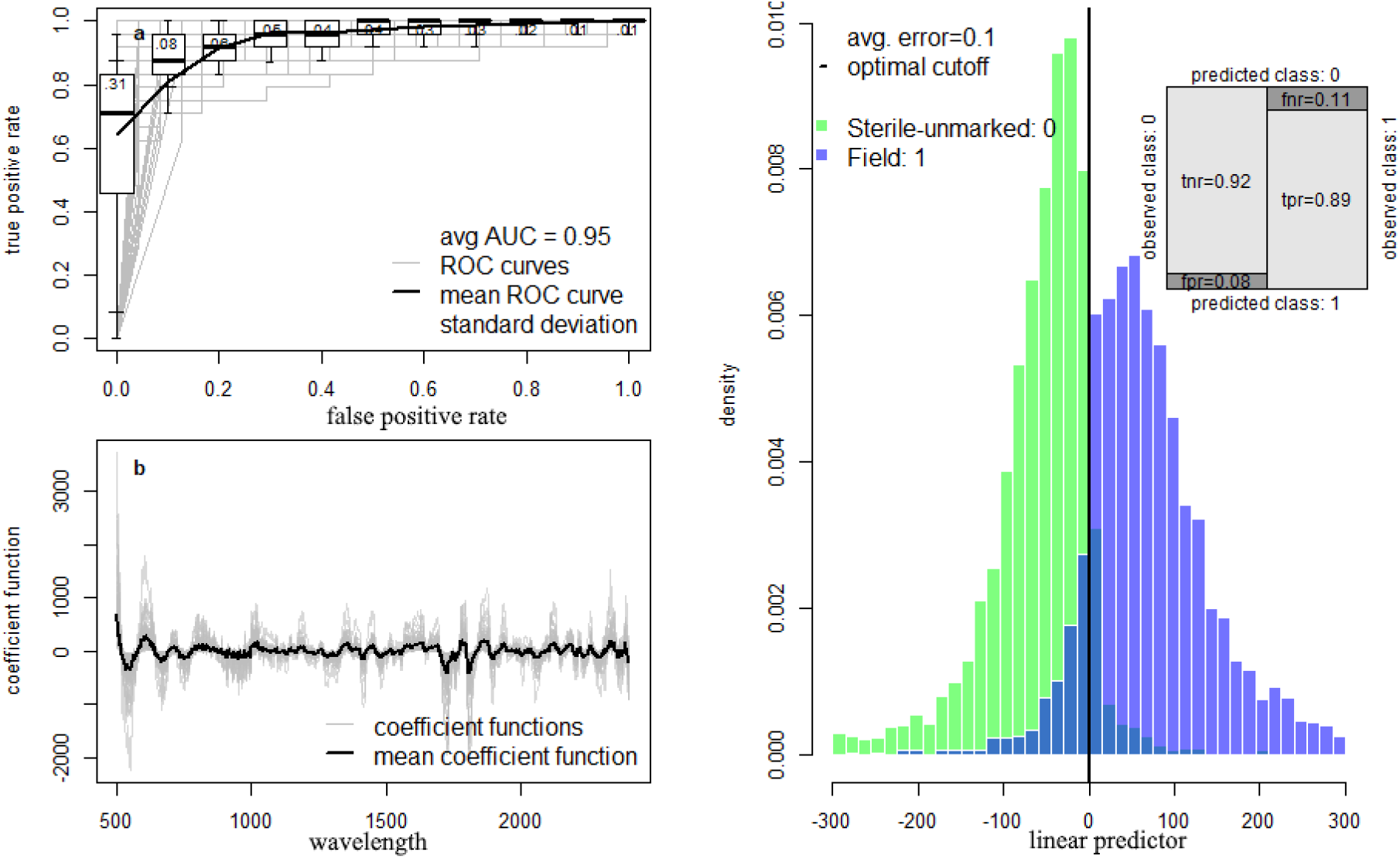
NIRS ability to discriminate between sterile unmarked and wild-caught flies. (a) Receiver operating characteristic (ROC) curve illustrating the diagnostic ability of the best-fit model. Overall performance is given by the average area under the ROC curve (AUC). A theoretical perfect diagnostic would be in the top left corner. The average ROC curve shown by the solid line with boxplots shows the variability for 100 randomizations of the training, validation and testing. (b) Coefficient functions for the best-fit model for each of the 100 dataset randomizations (grey lines) and the overall average (black line). (c) Histogram of the estimated linear predictor for the test observations, color-coded by the true class: marked irradiated flies (light blue colored bars) or Wild-caught flies (green bars). Vertical solid black line indicates the best threshold for differentiating marked irradiated flies and wild-caught flies. Darker blue bars indicate where the two distributions overlap and misclassified flies’ status. Misclassified wild-caught *Glossina* are shown to the left of the optimal classification threshold line and misclassified sterile unmarked *Glossina* to the right. Inset shows the confusion matrix illustrating the different error rates: true negative rate (tnr) for the sterile unmarked flies correctly classified; false negative rate (fnr) for the misclassified field flies; false positive rate (fpr) for the misclassified sterile unmarked flies; and true positive rate (tpr) corresponding to the field flies correctly classified.

However, using the group of sterile tsetse flies, NIRS was also able to distinguish between marked and unmarked flies with an accuracy of 81% (sterile-unmarked: 83%; sterile-marked: 79%; Fig 3a). The predictive models developed using spectra collected from laboratory unmarked and fertile versus laboratory unmarked sterile flies showed an accuracy of 88% (fertile-unmarked: 89%; sterile-unmarked: 87%; Fig 3b). Then, without any intervention operated during flies rearing in the insectary, NIRS was able to discriminate between unmarked fertile flies and wild-caught flies with 86% accuracy (fertile unmarked: 86%; wild-caught: 85%; Fig 3c). This highly accurate prediction of NIRS suggests that lab-reared flies naturally differ from their homologous wild type collected in the field.

**Figure 3:**
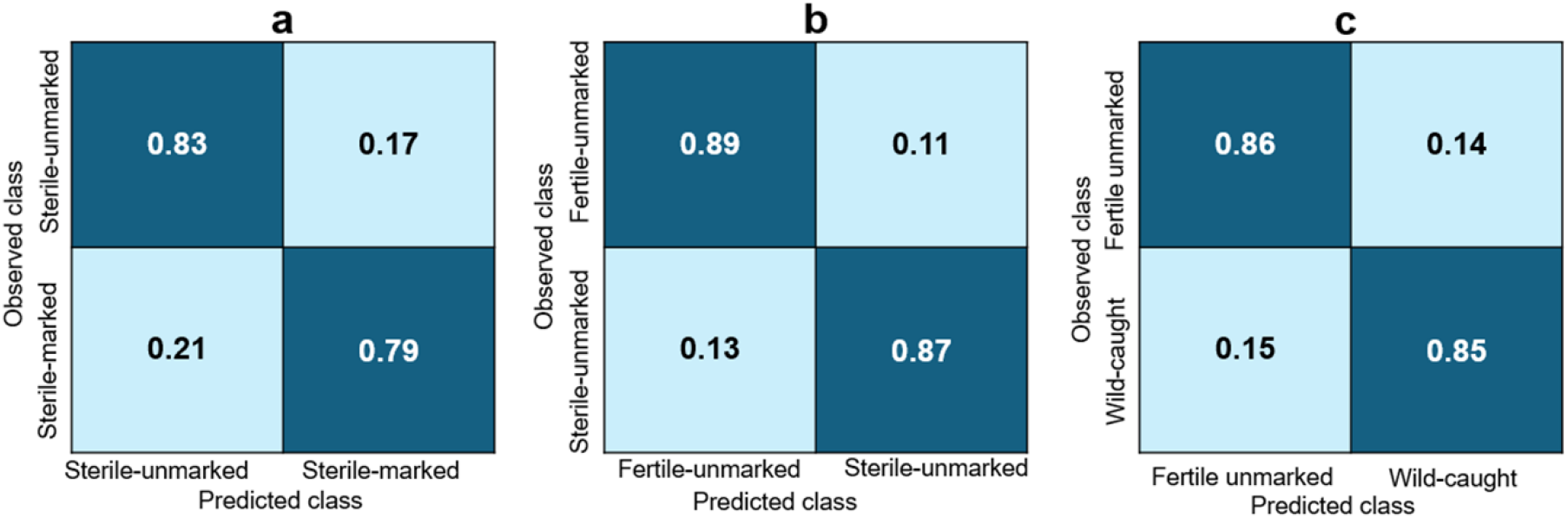
NIRS accuracy in predicting laboratory-reared and wild-caught *Glossina*. a: Marking effect (irradiated-unmarked *vs* irradiated-marked); b: Irradiation effect (unirradiated-unmarked *vs* irradiated-unmarked); c: Origin effect (laboratory fertile and unmarked *vs* wild).

Based on these results, the *Glossina* irradiation during the rearing (even without marking) could contribute to improving the NIRS ability to discriminate between marked sterile flies and wild-caught flies. This is interesting because the NIRS could be used as a complementary tool to discriminate doubtful flies (which have lost their marking) from the fluorescent camera.

## Discussion

The status of field-caught male tsetse (sterile or wild) is a crucial entomological parameter during an area-wide program that includes a sterile insect technique component. The fluorescence marking of sterile flies to differentiate them from wild ones is sometime defective to classify with the fluorescence camera. Filling these gaps leads to exploring the NIRS technique in the monitoring of trypanosomiasis vectors.

We showed for the first time that NIRS could be applied to accurately and rapidly discriminate between laboratory-reared sterile males from the wild-caught ones. This technique was able to differentiate with high accuracy (90%) the unmarked-sterile laboratory-reared flies from their wild-caught counterparts. The sterile males were not marked with fluorescent powder to better mimic the conditions of doubtful flies (having lost the fluorescent powder); therefore, they were unable for the fluorescent camera to differentiate them from wild males. The analysis scheme used in this study showed that the high predictive ability of NIRS would result from the combination of two distinctive features: the irradiation of laboratory-reared tsetse flies and the environmental living conditions. Indeed, the effect of irradiation was assessed between irradiated and non-irradiated laboratory males and NIRS was able to differentiate these two groups with 88% accuracy. The consequence of the irradiation on male flies does not only induce dominant lethal mutations in the gametes (LaChance, Schmidt, et Bushland 1967), but causes also somatic damage in the bodies of flies which leads to a reduction of the average life-span of irradiated males as compared with untreated ones (Vreysen, Van Der Vloedt, et Barnor 1996; Kaboré et al. 2023). These processes of deterioration of cellular contents caused by irradiation in flies could contribute to play a part in the differentiation between sterile lab flies and their wild counterparts by the NIRS.

On the other hand, the second scenario of explanatory analysis was carried out between the fertile laboratory and wild flies, where the NIRS technique discriminates them with an accuracy of 86%. This difference between laboratory and field flies could be due to environmental impacts (temperature, humidity, nutriment, etc.) on the tsetse fly structure and metabolism. Indeed, while laboratory flies are reared under controlled environmental conditions for over half a century, wild flies live in more natural setting conditions with important variations of several variables. This could lead to differences in body cellular composition between the two groups of flies, and contribute to this discrimination. Moreover, it has been shown that maintaining the same *G palpalis gambiensis* colony under artificial conditions for 43 years led to selection pressures which resulted in differences in sequences of the Cytochrome Oxydase I gene between the laboratory flies and their wild counterparts (Pagabeleguem et al. 2016).

In summary, the NIRS technique has an ability to differentiate interest groups of glossina with relatively good accuracy making it a promising method in the Xenomonitoring of trypanosomiasis control. Burkina Faso has been involved since 2006 in a Pan African program (PATTEC) that aimed at creating tsetse flies and trypanosomosis-free zones (Kabayo 2002). The first phase of PATTEC Burkina (2006-2013) managed to reduce fly densities by about 95 - 99% over an area of about 40,000 km2 (Percoma et al. 2018). PATTEC Burkina plans to define isolatable areas along the Mouhoun River where SIT could be applied. Moreover, the same isolated populations of *G. p. gambiensis* were identified in West Africa and might be targeted by eradication programs (Dicko et al. 2015). In terms of new methods of tsetse flies status determination, a molecular tool based on the mitochondrial gene COI (cytochrome oxidase I) has been developed and allowed to differentiate accurately wild tsetse males from sterile ones with good accuracy (Pagabeleguem et al. 2016). Although accurate, this method is costly, time-consuming for examining a large number of samples, destructive, laborious and requires well-trained personnel, therefore limiting its application in evaluating AW-IPM programs with a SIT component. In contrast, the NIRS technique could accurately and cost-effectively identify a larger number of trapped tsetse flies. Indeed, the initial cost of the instrument is approximately 40,000 USD, but there are no significant costs associated with running samples after this initial investment. The system can be battery powered and is field-portable. After calibrations are developed, the training required to use the system is only that required to operate a computer. Moreover, the NIRS technique becomes increasingly cost-effective as the number of samples for analysis increases (Mayagaya et al. 2009).

The NIRS technique, through its high accuracy and several advantages highlighted in many studies, appears important as a complementary tool when discrimination between sterile and wild tsetse flies with the fluorescence camera is doubtful. To the best of our knowledge, this is the first study to report the utility of the NIRS technique to accurately and rapidly determine the status (sterile or wild) or the origin (laboratory/field) of tsetse flies independently of the marking criteria. Based on these encouraging results, the NIRS technique could be optimized through further investigations to promote this new generation monitoring tool as a reliable long-term alternative for tsetse flies’ status/origin determination in the field and might be useful for other vector or insect pest control programs. The tsetse eradication program currently underway in the Niayes of Senegal (Seck et al. 2024) would be an opportunity to test this new method for validation.

## Notes

### Competing Interest Statement

The authors have declared no competing interest.

## References

Argilés-Herrero, Rafael, Gustavo Salvador-Herranz, Andrew G. Parker, Mario Zacarés, Assane G. Fall, Adji M. Gaye, Arooj Nawaz, Peter Takáč, Marc J. B. Vreysen, et Chantel J. de Beer. 2023. « Near-Infrared Imaging for Automated Tsetse Pupae Sex Sorting in Support of the Sterile Insect Technique ». Parasite 30:17. 10.1051/parasite/2023019.

Bauer, Burkhard, J Filledier, et Idrissa Kabore. 1984. « Large scale rearing of tsetse flies (Diptera, Glossinidae) in the C.R.T.A. Bobo-Dioulasso, Burkina based on in vitro feeding techniques ». Revue d’Elevage et de Medecine Veterinaire des Pays Tropicaux 37 (N° spécial): 9–17.

Bloem, S., J. Carpenter, A. Mccluskey, R. Fugger, S. Arthur, et S. Wood. 2007. « Suppression of the Codling Moth Cydia Pomonella in British Columbia, Canada Using an Area-Wide Integrated Approach with an SIT Components ». In Area-Wide Control of Insect Pests, édité par M. J. B. Vreysen, A. S. Robinson, et J. Hendrichs, 591–601. Dordrecht: Springer Netherlands. 10.1007/978-1-4020-6059-5_55.

Bouyer, J., M. T. Seck, et B. Sall. 2013. « Misleading Guidance for Decision Making on Tsetse Eradication: Response to Shaw et al. (2013) ». Preventive Veterinary Medicine 112 (3-4): 443–46. 10.1016/j.prevetmed.2013.05.017.

Challier, A, et C Laveissière. 1973. « Un nouveau piège pour la capture des glossines (Glossina: Diptera, Muscidae): description et essais sur le terrain ». Cahier O.R.S.T.O.M., série Entomologie médicale et Parasitologie 10 (4): 251–62.

Cuisance, D, H Politzar, P Mérot, et I Tamboura. 1984. « Les lâchers de mâles irradiés dans la campagne de lutte intégrée contre les glossines dans la zone pastorale de Sidéradougou, Burkina Faso ». Revue d’Elevage et de Médecine Vétérinaire des Pays Tropicaux 37 (4): 449–67.

Da, Dari F., Ruth McCabe, Bernard M. Somé, Pedro M. Esperança, Katarzyna A. Sala, Josua Blight, Andrew M. Blagborough, et al. 2021. « Detection of Plasmodium Falciparum in Laboratory-Reared and Naturally Infected Wild Mosquitoes Using near-Infrared Spectroscopy ». Scientific Reports 11. 10.1038/s41598-021-89715-1.

Depickère, Stéphanie, Antonio G. Ravelo-García, et Frédéric Lardeux. 2020. « Chagas Disease Vectors Identification Using Visible and Near-Infrared Spectroscopy ». Chemometrics and Intelligent Laboratory Systems 197 (février):103914. 10.1016/j.chemolab.2019.103914.

Dicko, Ahmadou H., Lassane Percoma, Adama Sow, Yahaya Adam, Charles Mahama, Issa Sidibé, Guiguigbaza-Kossigan Dayo, et al. 2015. « A Spatio-temporal Model of African Animal Trypanosomosis Risk ». PLoS Negl Trop Dis 9 (7): e0003921. 10.1371/journal.pntd.0003921.

Dowell, F.E., A.G. Parker, M.Q. Benedict, A.S. Robinson, A.B. Broce, et R.A. Wirtz. 2005. « Sex Separation of Tsetse Fly Pupae Using Near-Infrared Spectroscopy ». Bulletin of Entomological Research 95 (3): 249–57. 10.1079/BER2004357.

Esperança, Pedro M., Dari F. Da, Ben Lambert, Roch K. Dabiré, et Thomas S. Churcher. 2020. « Functional data analysis techniques to improve the generalizability of near-infrared spectral data for monitoring mosquito populations ». bioRxiv.

Goh, Brendon, Koek Ching, Ricardo J. Soares Magalhães, Silvia Ciocchetta, Michael D. Edstein, Rafael Maciel-de-Freitas, et Maggy T. Sikulu-Lord. 2021. « The Application of Spectroscopy Techniques for Diagnosis of Malaria Parasites and Arboviruses and Surveillance of Mosquito Vectors: A Systematic Review and Critical Appraisal of Evidence ». PLOS Neglected Tropical Diseases 15 (4): e0009218. 10.1371/journal.pntd.0009218.

Hendrichs, J., M. J. B. Vreysen, W. R. Enkerlin, et J. P. Cayol. 2021. « Strategic options in using sterile insects for area-wide integrated pest management ». In Sterile insect technique, 841–84. CRC Press.

Hursey, B. S., et J. Slingenbergh. 1995. « The tsetse fly and its effects on agriculture in sub-Saharan Africa ». World Animal Review, 67–73. http://link.springer.com/10.1007/1-4020-4051-2_22.

Itard, J. 1976. « L’élevage de Glossina palpalis gambiensis Vanderplank, 1949 (Diptera-Muscidae) à Maisons-Alfort ». Revue d’Elevage et de Medecine Veterinaire des Pays Tropicaux 29 (1): 43–58.

JN, Pollock. 1982. « Manuel de lutte contre la mouche tsé-tsé: Biologie, systématique et répartition des tsé-tsé ». Rome: FAO.

Kabayo, John P. 2002. « Aiming to eliminate tsetse from Africa ». Trends in parasitology 18 (11): 473–75.

Kaboré, Bénéwendé Aristide, Arooj Nawaj, Hamidou Maiga, Olga Soukia, Soumaïla Pagabeleguem, Marie Sophie Gisèle Ouédraogo/Sanon, Marc J. B. Vreysen, Robert L. Mach, et Chantel J. de Beer. 2023. « X-Rays Are as Effective as Gamma-Rays for the Sterilization of Glossina Palpalis Gambiensis Vanderplank, 1911 (Diptera: Glossinidae) for Use in the Sterile Insect Technique ». Scientific Reports 13 (1): 17633. 10.1038/s41598-023-44479-8.

Klassen, W., et C. F. Curtis. 2005. « History of the Sterile Insect Technique ». In Sterile Insect Technique, édité par V. A. Dyck, J. Hendrichs, et A.S. Robinson, 3–36. Berlin/Heidelberg: Springer-Verlag. 10.1007/1-4020-4051-2_1.

LaChance, L, C H Schmidt, et R C Bushland. 1967. « Radiation induced sterilization ». In Pest control: biological, physical and selected chemical methods., édité par W W Kilgore et R L Doutt, Academic, 147–96. New York, USA.

Laveissière, C., et P. Grébaut. 1990. « [The trapping of tsetse flies (Diptera: Glossinidae). Improvement of a model: the Vavoua trap] ». Tropical medicine and parasitology: official organ of Deutsche Tropenmedizinische Gesellschaft and of Deutsche Gesellschaft fur Technische Zusammenarbeit (GTZ) 41 (2): 185–92.

Maia, Marta F., Melissa Kapulu, Michelle Muthui, Martin G. Wagah, Heather M. Ferguson, Floyd E. Dowell, Francesco Baldini, et Lisa Ranford-Cartwright. 2019. « Detection of Plasmodium Falciparum Infected Anopheles Gambiae Using Near-Infrared Spectroscopy ». Malaria Journal 18 (1): 85. 10.1186/s12936-019-2719-9.

Mayagaya, Valeliana S., Kristin Michel, Mark Q. Benedict, Gerry F. Killeen, Robert A. Wirtz, Heather M. Ferguson, et Floyd E. Dowell. 2009. « Non-Destructive Determination of Age and Species of Anopheles Gambiae s.l. Using near-Infrared Spectroscopy ». The American Journal of Tropical Medicine and Hygiene 81 (4): 622–30. 10.4269/ajtmh.2009.09-0192.

Musiime, Alex K., Joseph Okoth, Melissa Conrad, Daniel Ayo, Ismail Onyige, John Rek, Joaniter I. Nankabirwa, Emmanuel Arinaitwe, Moses R. Kamya, et Grant Dorsey. 2019. « Is that a real oocyst? Insectary establishment and identification of Plasmodium falciparum oocysts in midguts of Anopheles mosquitoes fed on infected human blood in Tororo, Uganda ». Malaria journal 18 (1): 1–11.

Pagabeleguem, Soumaïla, Geoffrey Gimonneau, Momar Talla Seck, Marc J. B. Vreysen, Baba Sall, Jean-Baptiste Rayaissé, Issa Sidibé, Jérémy Bouyer, et Sophie Ravel. 2016. « A Molecular Method to Discriminate between Mass-Reared Sterile and Wild Tsetse Flies during Eradication Programmes That Have a Sterile Insect Technique Component ». PLOS Neglected Tropical Diseases 10 (2): e0004491. 10.1371/journal.pntd.0004491.

Pagabeleguem, Soumaïla, Oumar Koughuindida, Ernest Wendemanegde Salou, Geoffrey Gimonneau, Ange Irénée Toé, Bénéwendé Aristide Kaboré, Kiswend-sida Mikhailou Dera, et al. 2023. « Gamma-Radiation of Glossina Palpalis Gambiensis Revisited: Effect on Fertility and Mating Competitiveness ». Parasite 30:8. 10.1051/parasite/2023009.

Pagabeleguem, Soumaïla, Momar T. Seck, Baba Sall, Marc J.B. Vreysen, Geoffrey Gimonneau, Assane G. Fall, Mireille Bassene, et al. 2015. « Long Distance Transport of Irradiated Male Glossina Palpalis Gambiensis Pupae and Its Impact on Sterile Male Yield ». Parasites & Vectors 8 (1): 259. 10.1186/s13071-015-0869-3.

Pagabeleguem, Soumaïla, Ange Irénée Toé, Sié Hermann Pooda, Kiswendsida Mikhailou Dera, Abdou Salam Belem, Adrien Marie Gaston Belem, Gisèle Marie Sophie Ouedraogo/Sanou, et al. 2021. « Optimizing the feeding frequency to maximize the production of sterile males in tsetse mass-rearing colonies ». PLOS ONE 16 (1): e0245503. 10.1371/journal.pone.0245503.

Percoma, Lassané, Adama Sow, Soumaïla Pagabeleguem, Ahmadou H. Dicko, Oumarou Serdebéogo, Mariam Ouédraogo, Jean-Baptiste Rayaissé, Jérémy Bouyer, Adrien M. G. Belem, et Issa Sidibé. 2018. « Impact of an integrated control campaign on tsetse populations in Burkina Faso ». Parasites & Vectors 11 (1): 270. 10.1186/s13071-017-2609-3.

Politzar, H, et D Cuisance. 1984. « An integrated campaign against riverine tsetse flies Glossina palpalis gambiensis and Glossina tachinoides by trapping and the release of sterile males. » Insect science and application 5:439–42.

Seck, Momar T., Soumaïla Pagabeleguem, Mireille Bassene, Assane G. Fall, Thérèse A.R. Diouf, Baba Sall, Marc J.B. Vreysen, et al. 2015. « Quality of sterile male Glossina palpalis gambiensis tsetse after long distance transport as chilled, irradiated pupae ». PLoS Neglected Tropical Diseases 9 (11): e0004229. 10.1371/journal.pntd.0004229.

Seck, Momar Talla, Assane Guèye Fall, Mamadou Ciss, Mame Thierno Bakhoum, Baba Sall, Adji Marème Gaye, Geoffrey Gimonneau, et al. 2024. « Animal Trypanosomosis Eliminated in a Major Livestock Production Region in Senegal Following the Eradication of a Tsetse Population ». Parasite 31:11. 10.1051/parasite/2024010.

Sikulu, Maggy, Gerry F. Killeen, Leon E. Hugo, Peter A. Ryan, Kayla M. Dowell, Robert A. Wirtz, Sarah J. Moore, et Floyd E. Dowell. 2010. « Near-Infrared Spectroscopy as a Complementary Age Grading and Species Identification Tool for African Malaria Vectors ». Parasites & Vectors 3 (juin):49. 10.1186/1756-3305-3-49.

Sikulu-Lord, Maggy T., Michael D. Edstein, Brendon Goh, Anton R. Lord, Jye A. Travis, Floyd E. Dowell, Geoffrey W. Birrell, et Marina Chavchich. 2024. « Rapid and Non-Invasive Detection of Malaria Parasites Using near-Infrared Spectroscopy and Machine Learning ». PLOS ONE 19 (3): e0289232. 10.1371/journal.pone.0289232.

Swallow, Brent. 1999. « Impacts of Trypanosomosis on African Agriculture ». Rome, Italy.

Takken, V., M. A. Oladunmade, L. Dengwat, H. U. Feldmann, J. A. Onah, S. O. Tenabe, et H. J. Hamann. 1986. « The eradication of Glossina palpalis palpalis (Robineau-Desvoidy)(Diptera: Glossinidae) using traps, insecticide-impregnated targets and the sterile insect technique in central Nigeria ». Bulletin of Entomological Research 76 (2): 275–86.

Tátila-Ferreira, Aline, Gabriela A. Garcia, Lilha dos Santos, Márcio G. Pavan, Carlos José de C Moreira, Juliana C. Victoriano, Renato da Silva-Junior, Jacenir R. dos Santos-Mallet, Thaiane Verly, et Constança Britto. 2021. « Near infrared spectroscopy accurately detects Trypanosoma cruzi non-destructively in midguts, rectum and excreta samples of Triatoma infestans ». Scientific reports 11 (1): 1–10.

Valladares, Vanessa, Célio Pasquini, Silvana C. Thiengo, Monica A. Fernandez, et Clélia C. Mello-Silva. 2021. « Field Application of NIR Spectroscopy for the Discrimination of the Biomphalaria Species That Are Intermediate Hosts of Schistosoma Mansoni in Brazil ». Frontiers in Public Health 9:636206. 10.3389/fpubh.2021.636206.

Vargas-Terán, M., B. S. Hursey, et E. P. Cunningham. 1994. « Eradication of the Screwworm from Libya Using the Sterile Insect Technique ». Parasitology Today (Personal Ed.) 10 (3): 119–22. 10.1016/0169-4758(94)90014-0.

Vreysen, M. J., K. M. Saleh, M. Y. Ali, A. M. Abdulla, Z. R. Zhu, K. G. Juma, V. A. Dyck, A. R. Msangi, P. A. Mkonyi, et H. U. Feldmann. 2000. « Glossina Austeni (Diptera: Glossinidae) Eradicated on the Island of Unguja, Zanzibar, Using the Sterile Insect Technique ». Journal of Economic Entomology 93 (1): 123–35. 10.1603/0022-0493-93.1.123.

Vreysen, Marc J.B. 2005. « Monitoring Sterile and Wild Insects in Area-Wide Integrated Pest Management Programmes ». In Sterile Insect Technique, édité par Victor A. Dyck, J. Hendrichs, et Alan S. Robinson, 325-61. Vienna, Austria: Springer Netherlands. http://link.springer.com/chapter/10.1007/1-4020-4051-2_12.

Vreysen, Marc J.B., Khalfan M. Saleh, Mashavu Y. Ali, Abdulla M. Abdulla, Zeng-Rong Zhu, Kassim G. Juma, Arnold V. Dyck, Atway R. Msangi, Paul A. Mkonyi, et Udo Feldmann. 2000. « Glossina Austeni (Diptera: Glossinidae) Eradicated on the Island of Unguja, Zanzibar, Using the Sterile Insect Technique ». Journal of Economic Entomology 93 (1): 123–35. 10.1603/0022-0493-93.1.123.

Vreysen, Marc J.B., Khalfan M. Saleh, Furaha Mramba, Andrew G. Parker, Udo Feldmann, Victor A. Dyck, Atway Msangi, et Jérémy Bouyer. 2014. « Sterile insects to enhance agricultural development: the case of sustainable tsetse eradication on Unguja Island, Zanzibar, using an area-wide integrated pest management approach ». PLoS neglected tropical diseases 8 (5): e2857.

Vreysen, Marc J.B., M. Seck, Baba Sall, Abdou Mbaye, M. Bassene, Assane Fall, M. Lo, et Jérémy Bouyer. 2021. « Area-Wide Integrated Management of a Glossina palpalis gambiensis Population from the Niayes Area of Senegal: A Review of Operational Research in Support of a Phased Conditional Approach ». In Area-Wide Integrated Pest Management: Development and Field Application, par J. Hendrichs, R. Pereira, et M. J. B. Vreysen, 275–303. Boca Raton, Florida, USA: CRC Press.

Vreysen, Marc J.B., Momar Talla Seck, Baba Sall, et Jérémy Bouyer. 2013. « Tsetse Flies: Their Biology and Control Using Area-Wide Integrated Pest Management Approaches ». Journal of Invertebrate Pathology 112 (mars):S15-25. 10.1016/j.jip.2012.07.026.

Vreysen, Marc J.B., A. Van Der Vloedt, et H. Barnor. 1996. « Comparative Gamma-Radiation Sensitivity of Glossina Tachinoides Westv., Glossina Fuscipes Fuscipes Newst. and Glossina Brevipalpis Newst. (Diptera, Glossinidae) ». International Journal of Radiation Biology 69 (1): 67–74.

